# In Vitro and Insilico screening platform for the identification of aldose reductase inhibitors for antidiabetic lead compounds from *Abutilon indicum (L.)*

**DOI:** 10.1101/2020.10.31.363549

**Authors:** L Lavanya, V. Veeraraghavan, CN Prashantha, Renuka Srihari

**Author notes:** **Corresponding Author:** V.Veeraraghavan, Professor, Department of Biochemistry, School of Applied Sciences, REVA University, Bengaluru-560064., **Email:**.

## Abstract

**Introduction:** Type 2 Diabetes Mellitus (T2DM) is a long-term metabolic disorder that primarily characterized by impaired insulin resistance to become hyperglycemia. People suffering from T2DM have a higher risk of developing various diseases but, on top of that, some diabetic drugs are also suspected of increasing the risk in some cases. Aldose reductase is a key target enzyme to catalyze the reduction of glucose to sorbitol and does not readily diffuse across cell membranes and cause retinopathy and neuropathy. The aldolase reductase inhibitors prevent the conversion of glucose to sorbitol and may have the capacity of preventing and / or treating several diabetic complications. It will be expected to be twofold in the subsequent decade due to intensification in the senile population with the number of people affected, thus adding to the liability on medical providers in poor developed countries using herbal medicine to control the diabetes. In recent investigation, the antidiabetic property of phytochemicals extracted from leafs of *Abutilon indicum (L.)* is elucidated using animal models.

**Materials and Methods:** In the current study using aldose reductase enzyme assay inhibitor of Rat lens Aldose reductase were treated with *A. indicum* methanolic leaf extract at different concentrations (6.25, 12.5, 25, 50, 100, and 200μg/mL). Copper sulphate was used as reference drug and docking studies to predict the screen the best aldose reductase inhibitor.

**Results and Discussion:** The crude extract exhibited cytotoxicity against rat lens aldose reductase (IC_50_ = 135.8 μg/L vs ref 13.60 μg/L) using In Vitro. The docking is performed with 11 compounds shows Ertugliflozin, 9H-Cycloisolongifolene, 8-oxo and 7-hydroxy cadalene showed a good binding interaction with aldose reductase.

**Conclusion:** We are concluding that the invitro and *in silico* analysis helps researchers to utilize these compound for aldose reductase inhibitors and further can be used for clinical applications.

## 1.0 Introduction

Diabetes mellitus is an endocrine disorder in which glucose metabolism is impaired because of total loss of insulin secretion and mechanism of action after destruction of pancreatic β-cells or insensitivity of target tissues to insulin in non-insulin dependent diabetes mellitus ^[1–3]^. People suffering from T2DM have a higher risk of developing various diseases such as diabetic neuropathy ^[4]^ and retinopathy ^[5]^. With the theme of T2DM, over 26.2 million people were observed in India by 2019 ^[6]^ and its alarming mark of 69.9 million by 2025 ^[7]^. A serious issue in diabetic patients with the complications of diabetic mellitus with various diseases ^[8]^. The excessive amount of glucose circulating in blood plasma cause asymptomatic glucose metabolism ^[9]^. In general the glucose level to the body should range from ~5.6 and ~7 mmol/l (100–126 mg/dl) and this range is increased to ~11.1 mmol/l (200 mg/dl) but there is no much of the problem in the tissues or organs, but this range get increase to 15–20 mmol/l (~250–300 mg/dl) this results in hyperglycemia ^[10]^. Many theories have been explained that glucose is converting to fructose via hexokinase in glycolysis pathway. Over 33% of excess glucose is involved in polyol pathway ^[11–13]^.

The polyol pathway of glucose metabolism is stimulated under the excess of glucose condition in the blood plasma that converts glucose to sorbitol by utilizing aldose reductase with the reduced oxidation of NADPH to NADP+. However, the sorbitol is converting to fructose by utilizing sorbitol dehydrogenase with the oxidation of NADH to NAD+ under normoglycemic conditions ^[14–15]^. The uncontrolled high blood glucose increases the activation of aldose reductase in cytosolic region that catalyzes carbonyl-containing compounds such as aldehydes, ketones, monosaccharaides etc ^[16–17]^. It also reduces low enzymatic activity with all-trans-retinal, 9-cis-retinal, and 13-cis-retinal ^[18]^. Based on the study, aldose reductase plays an important role to complicate diabetes ^[19,20]^. Type 2 diabetes increases the availability of glucose in insulin – insensitive tissues such as lens, nerve and retina leads to the increased formation of sorbitol through the polyol pathway ^[21]^. Sorbitol does not readily diffuse across cell membranes and the intra cellular accumulation of sorbitol has been implicated in the chronic complications of diabetes such as cataract, neuropathy and retinopathy ^[22]^. There are several mutations observed in the aldose reductase protein sequence with the region of Asp44➔ Asn44, Lys78 ➔ Met78 and His111➔ Asn111 mutations that reduce glyceraldehydes oxidoreductase activity that shows high risk of diabetic retinopathy with the subgroup of diabetes mellitus ^[23–24]^. These findings suggest that Aldose reductase inhibitors prevent the conversion of glucose to sorbitol and may have the capacity of preventing and / or treating several diabetic complications ^[25]^.

There are several treatment methods helps to aldose reductase inhibition that convert glucose to sorbitol, but still there is a lot of demand to predict the alternate treatment methods by utilizing natural products for the inhibition of aldose reductase for the treatment of diabetic nephropathy and retinopathy. In the current research is used *Abutilon indicum (L.)* Sweet has long history of antidiabetic properties, the plant parts such as leaf, stem and roots have rich source of flavonoids, phenolics compounds, steroids, alkaloids, tannins, saponins and glycosides ^[26]^.The powdered sample of A.indicum leaf sample have rich source of medicinal properties for the treatment of various disease such as anti-inflammatory, anthelmintic, analgesic and antidiabetic properties. The rhizoids can use for the treatment of leprosy, ulcers, headaches, gonorrhea, and bladder infection. Based on the literature, administration of *A. indicum* extract at 0.25 and 0.5 g kg−1 resulted in significant decreases of postprandial plasma blood glucose. The in diabetic rat when compared with diabetic control rats (p < 0.05) ^[27]^.

The aim of the present study is to identify the phytochemicals present in *A.indicum* leaf extract to assess aldose reductase inhibition using in vivo study and insilico docking study to screen the phytochemicals as potential aldose reductase inhibitors.

## 2.0. Materials and Methods

### 2.1. Identification of phytochemicals

A fresh and health leaves of *A.indicum* was collected and extracted using standard protocol ^(7)^. The spectroscopic analysis is performed using GC-MS in Bangalore analytical research center, Bangalore ^[28]^. The gas chromatography with the mode of Trace-1310 GC coupled to Triple quad mass spectrometer (MS) of the model TSQ8000 from Thermo Scientific is used to perform the analysis. The TGSMS column with dimension of 30m×0.25mm×0.25um was used for the analysis. 1.0μL of clear extract was injected into GC-MS with an oven temperature at 40°C with hold time of 2 min with the carrier gas flow rate of 1 ml/min up to 5 min to 280°C and hold time of 10 min at the flow rate of 20°C/min. The split sampling technique was used to inject the sample in the ratio of 1:30, Retention indices (RI) of the compounds were determined by comparing the retention times of a series and identification of each component was confirmed by comparison of its retention index with mass is from 50 to 600 kDa in a full scan. The carrier gas used in the analysis was helium which had the flow rate of 1ml/min. Interpretation of Mass-Spectrum was carried out by using the database of National institute Standard and Technology (NIST) having more than 62,000 patterns. The spectrum of the unknown components was compared with the spectrum of known components which was stored in the NIST library. The molecular weight, name, chemical structure and molecular formula of the components of the test materials were ascertained. The peak in GCMS of methanol extract of leaf of *A.indicum* showed the presence of the secondary phytochemical compounds like phenolics and fatty acids and its esters.

### 2.2. Antidiabetic assay analysis

#### 2.2.1 Rat lens Aldose reductase preparation

Crude Aldose reductase was prepared from rat lens. Eyeballs were removed from 9 week old male rats. Animal care and protocols were in accordance with and approved by Institutional Animal Ethics Committee. Lenses were dissected by posterior approach and homogenized in 10 volumes of 100 mM potassium phosphate buffer pH 6.2. The homogenate was centrifuged at 15,000 Xg for 30 min at 4°C and the resulting supernatant was used as the source of Aldose reductase ^[31]^. The assay mixture in 1 mL contained 50 μM potassium phosphate buffer pH 6.2, 0.4 mM lithium sulfate, 5 μM 2-mercaptoethanol, 10 μM DL-glyceraldehydes, 0.1 μM NADPH and enzyme preparation (rat lens enzyme). Appropriate blanks were employed for corrections. The assay mixture was incubated at 37°C and initiated by the addition of NADPH at 37°C. The change in the absorbance at 340 nm due to NADPH oxidation was measured spectrophotometrically.

### 2.3. Antioxidant Analysis of Aldose reductase enzyme

#### 2.3.1. ABTS assay Analysis

ABTS assay is based on the scavenging of light by ABTS radicals. An antioxidant with an ability to donate a hydrogen atom will quench the stable free radical, a process which is associated with a change in absorption which can be followed spectroscopically. The relatively stable ABTS radical has a green color and is quantified spectrophotometrically at 734nm ^[29,30]^.

##### PBS (Phosphate Buffered Saline

125mM NaCl in 10mM Sodium phosphate buffer, pH 7.4):0.14196g of Disodium hydrogen orthophosphate, 0.1560g of Sodium dehydrogenase orthophosphate and 0.7305g of Sodium chloride is dissolved in 25ml of distilled water and pH is adjusted to 7.4 using dilute sodium hydroxide solution and the volume is made up to 100ml with de-ionized water.

##### ABTS (2, 2’-azinobis-ethyl-benzothiozoline-6-sulphonic acid) (7mM)

38.4mg of ABTS is dissolved in PBS and the volume is made up to 10ml.

##### APS (Ammonium per sulfate) (2.45mM)

5.59mg of APS is dissolved in PBS and the volume made up to 10ml.

##### ABTS (2, 2’-azinobis-ethyl-benzothiozoline-6-sulphonic acid) Radical solution

a. **Mother stock:** 10ml of ABTS (7mM) and 10ml of APS (2.45mM) solutions are mixed and allowed to maintain at room temperature in dark for 16 hours.
b. **Working solution:** 1.7ml of the mother stock is made up to 50ml with PBS, so as to give an absorbance of 1.000

##### Quercetin (1000μg/mL) (Reference standard)

1mg was made up to 1mL with PBS. The assay is performed as per Auddy (2003) ^[32]^. ABTS radical cations are produced by reacting ABTS and APS on incubating the mixture at room temperature in dark for 16 hours. The solution thus obtained is further diluted with PBS to give an absorbance of 1.000. Different concentrations of the test sample and the reference standard (highest volume taken was 50μl) are added to 950μL of ABTS working solution to give a final volume of 1ml, made up by adding PBS. The absorbance is recorded immediately at 734nm. The percent inhibition is calculated at different concentrations and the IC50 values are calculated using Graph pad prism software.

Calculating percentage growth inhibition:

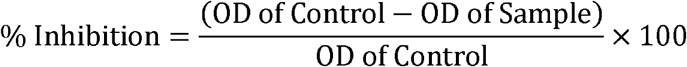

### 2.4. Statistical Analysis

The data were presented as the mean ± standard error of the mean (SEM). Statistical analysis was performed using the Student’s t-test or analysis of variance (ANOVA) followed by the least significant differences (LSD) test. The significance of differences was set at P-value < 0.05.

### 2.5. Computational Analysis of phytochemicals and screening

Using GC-MS analysis, we have predicted 8 chemical compounds such as Undecane, Benzeneethanmine, Astaxanthin, Diosgenin, Vildagliptin, Isospathulenol, Ertugliflozin, 9H-Cycloisolongifolene,8-oxo, 7-Hydroxycadalene, Hentriacontane, and Tetratetracontane compounds was designed using ACD/Chemsketch 11.0. Pharmacophore analysis is performed to understand the drug like character of the chemicals based on Lipinski Rule of 5 with selected parameters such as logP, TPSA, molecular weight, hydrogen bond donor, hydrogen bond acceptor, volume, number of rotatable bonds and total number of atoms using molinspiration online tool to calculate the molecular property and bioactive scores such as GPCR, ion channel, kinase inhibitor, nuclear receptor, protease inhibitor and enzyme inhibitor properties. This result helps to understand the functional inhibition of phytochemical against aldose reductase inhibitor. ADMET analysis is performed to the selected compounds using admetSAR tool to screen the compounds based on absorption, distribution, metabolism, excretion and toxicity prediction. The most important parameters such as blood brain barer, acute toxicity, carcinogenicity, LD_50_, maximum recommended daily dose and Mutagenicity is predicted by lazar toxicity predictions server.

### 2.6. Protein modeling and Docking

The aldose reductase enzyme structure id retrieved using Protein Data Bank (PDB) 1ABN. The selected protein structure is modeled and validated using Swiss PDB viewer to understand the complexity of the protein structure, ramachandran plot is predicted to understand the accuracy of the predicted protein model using RAMPAGE server. A classic ramachandran plot of Ψ versus Φ conformational angels of 3D macromolecule measures the torsion angels of Cα (ideal)-N-C_β_ (obs). Active site amino acids were predicted using CastP calculation server based on the delineating measures of surface regions such as surface area and surface volume of 3D protein structure. Docking analysis of aldose reductase and 11 phytochemicals extracted from A.indicum

(L) along with reference drug compound Diosgenin, Vildagliptin, Ertugliflozin and Quercetin was performed using AutoDock v.4.2, wherein we added Gasteiger chargers and hydrogen atoms to the polar group of amino acids the macromolecule, whereas ligand structure by adding torsion counts of amide bonds rotatable and all active bonds non-rotatable and generated the PDBQT file for both protein and ligand structures. Grid space was set in Autogrid by selecting important residues Gly18, Tyr209, Ser210, Leu212, Ser214, Pro261, Lys262 and Ser263 with the grid box size *x=60 Å, y=60 Å* and *z=60 Å* and grid spacing of 0.375 Å that provides search space, the grid center was selected at dimensions *x=−0.056, y=0.028, z=−0.083* used to calculate grid parameters that helps to understand the grid energy with equilibrated energy distribution. AutoDock was used to dock aldose reductase with chemical compounds by adding AutoDock parameters such as Lamarckian genetic algorithm (LGA) with default parameters. The best docking confirmation of protein ligand interactions is predicted with the energy value in kcal mol−1 and the followed by the analysis of hydrogen bonding interaction and the hydrophobic interaction. The best docking complex of aldose reductase-ligand was selected and used as an input for molecular dynamic simulations.

## 3 3.0 Results

### 3.1. Plant extract and GC-MS Analysis

The extraction of the phytochemicals from *A.indicum* was performed using methanol extraction method using standard protocol and the resultant fractions were analyzed using GC-MS (figure: 1). The GC-MS is performed to identify the constituents of volatile matter, long chain, branched chain hydrocarbons, and glycosides were identified based on peak area, retention time and molecular weight to confirm phytochemicals. Based on the NIST library comparison, we predicted eight compounds which revealed the presence of chemical structures (table: 1). the selected phytochemicals were screened to predict antidiabetic property for aldose reductase inhibitor prediction.

**Figure: 1.**
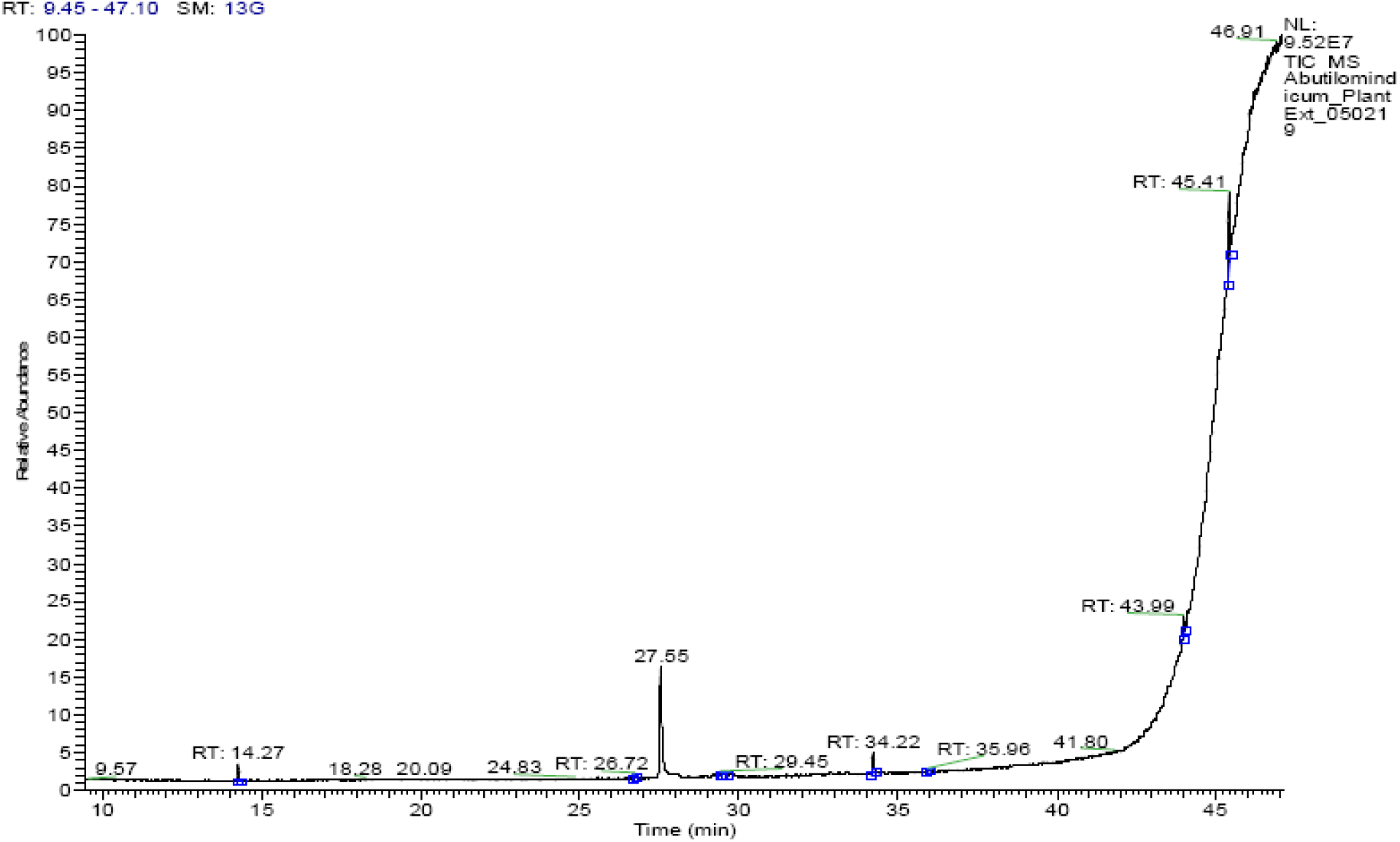
GC-MS analysis of *A.indicum* leaf extract.

**Table: 1.** Bioactivity prediction of phytochemicals isolated from *A.indicum* leaf extract.

| S.No | Retention time | Peak Area % | Name of the compound | Molecular Formula | Molecular Weight (g/mol) |
| --- | --- | --- | --- | --- | --- |
| 1 | 14.27 | 13.51 | Undecane | C<sub>11</sub>H<sub>24</sub> | 156.313 |
| 2 | 26.72 | 2.3 | Benzeneethanmine, 2,5-dimethoxy-α,4-dimethyl- | C<sub>8</sub>H<sub>11</sub>N | 121.18 |
| 3 | 29.45 | 2.04 | Astaxanthin | C<sub>40</sub>H<sub>52</sub>O<sub>4</sub> | 596.8 |
| 4 | 29.54 | 2.65 | Isospathulenol | C<sub>15</sub>H<sub>24</sub>O | 220.35 |
| 5 | 34.22 | 26.3 | 9H-Cycloisolongifolene, 8-oxo- | C<sub>15</sub>H<sub>22</sub>O | 218.33 |
| 6 | 35.96 | 2.09 | 7-Hydroxycadalene | C<sub>15</sub>H<sub>18</sub>O | 214.3 |
| 7 | 43.99 | 9.59 | Hentriacontane | C<sub>31</sub>H<sub>64</sub> | 436.8 |
| 8 | 45.41 | 41.51 | Tetratetracontane | C<sub>44</sub>H<sub>90</sub> | 619.2 |

### 3.2. Aldose reductase inhibitory activity prediction

The selected aldose reductase inhibitory assay activity was tested using copper sulphate as a substrate along with the fractions of phytochemicals with concentration of 0, 2.5, 3.125, 6.25, 12.5, 25, and 50μg/ml. The aldose reductase inhibition rate was calculated as percentage with respect to the control value and expressed as mean of standard and tested samples. The concentration of each test sample giving rise to 50% inhibition of activity (IC_50_) was estimated from the least-squares regression line of the logarithmic concentration plotted against inhibitory activity. Copper sulphate was used as positive control. The results shows IC_50_= 13.60 μg/l of copper sulphate along with IC_50_= 135.8 μg/l of phytochemical extract as best aldose reductase inhibitors (Figure: 2a; 2b).

**Figure: 2a.**
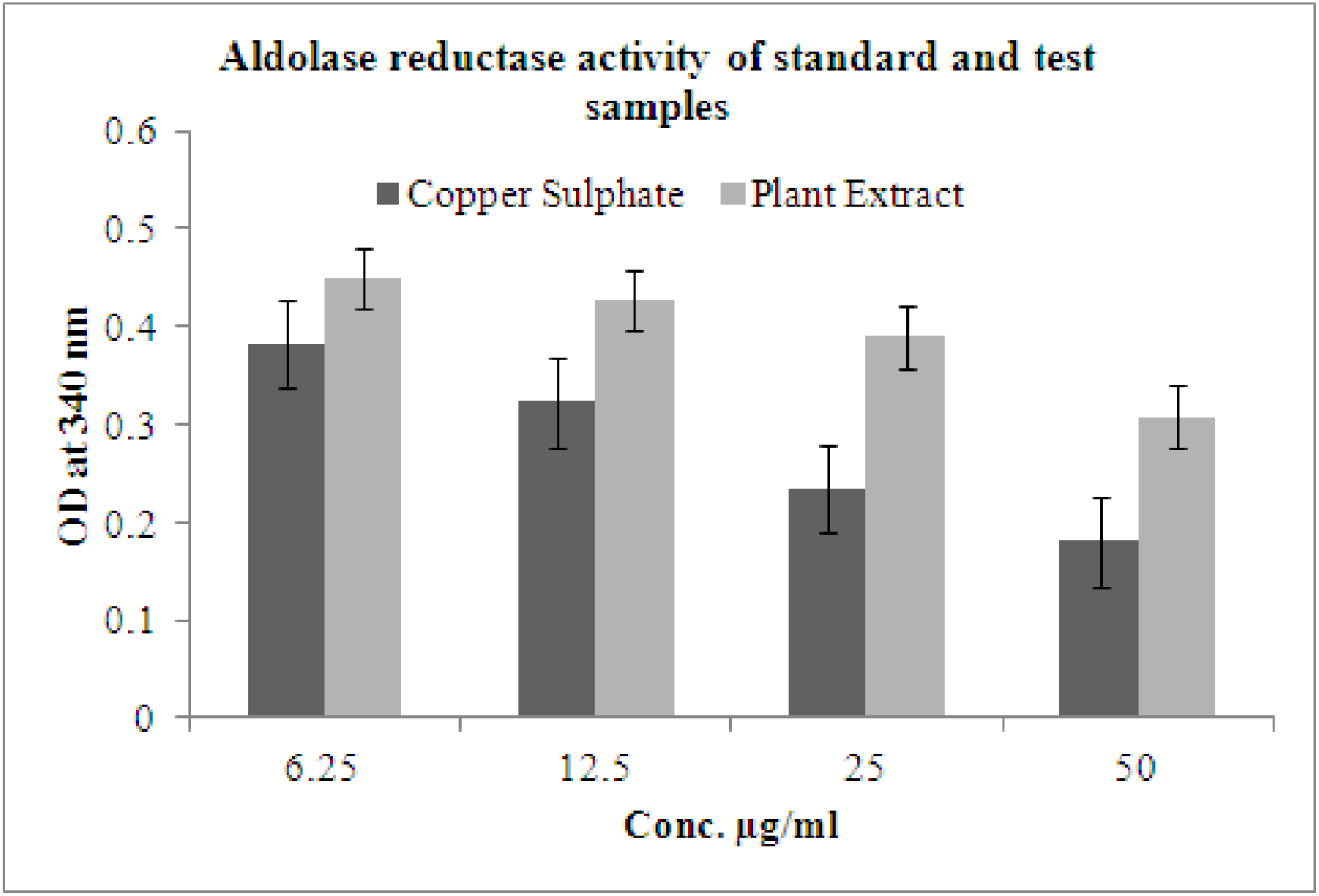
Spectroscopic analysis of aldose reductase inhibitory study using standard and test samples.

**Figure: 2b.**
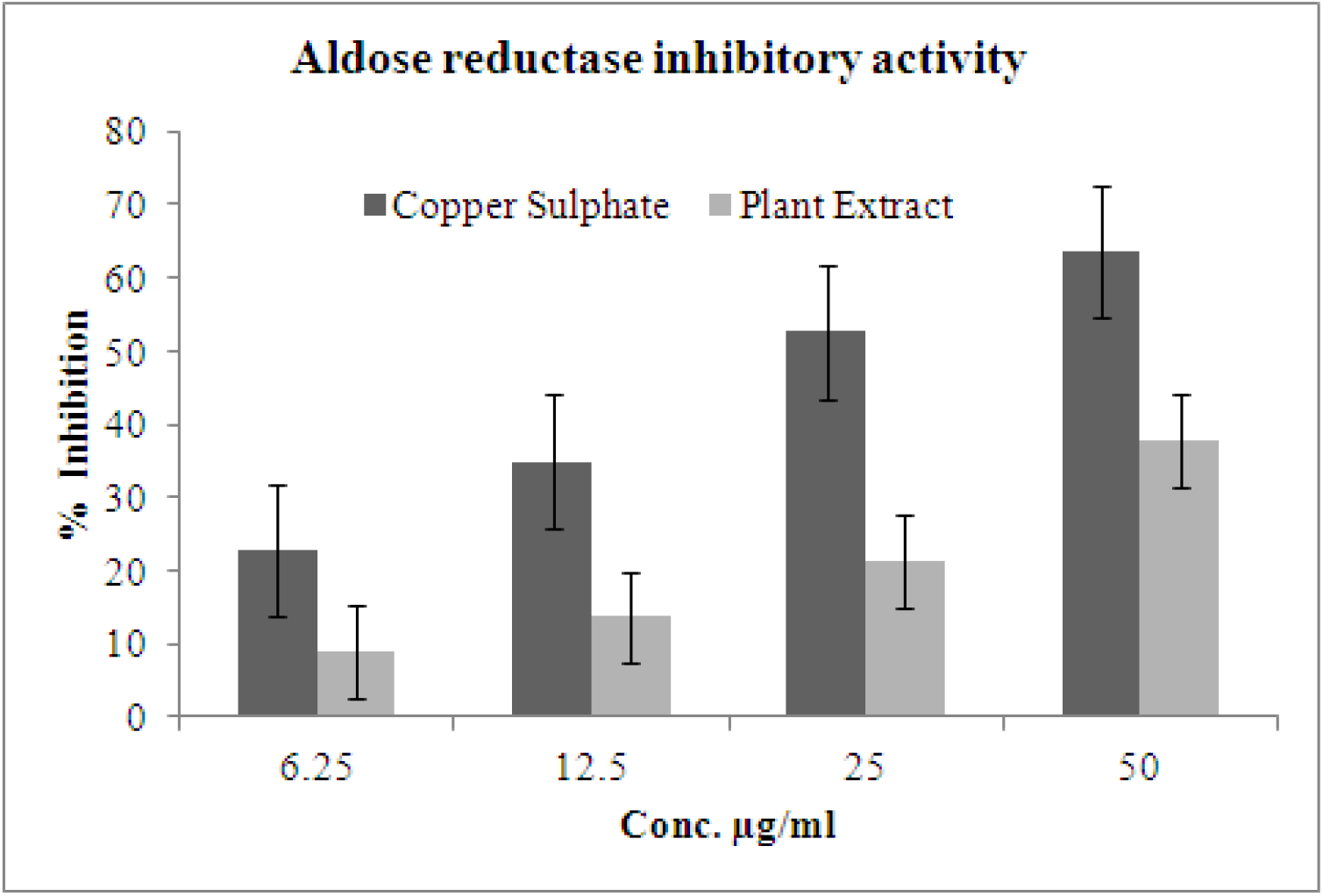
Percentage of inhibition were predicted with aldose reductase with standard and test samples.

### 3.3. Antioxidant analysis of ABTS assays

The ABTS assay of free radical scavenging assays are widely used for antioxidant activity determination of various plant extracts. Here, we used *A.indicum* leaf extract is used to predict based on flavonoids to scavenge ABTS+ radical. Antioxidant activity was expressed in terms of percentage inhibition for different extracts and a graph was plotted between percentage inhibition and concentration of different extracts. Antioxidant activity showed an increase with the increase in concentration. The IC50 value of methanol extract was lowest as compared to other extracts. IC50 of extracts were ranged from LogIC_50_= 0.08876μg/ml and IC_50_= 1.227μg/ml with Quercetin structure and phytochemicals having the LogIC50= 1.850μg/ml and IC_50_= 70.78μg/ml (Figure: 3a, 3b). Our results have proved that methanol extract of *A.indicum* have significant free radical scavenging activity and therefore exhibited higher antioxidant activities.

**Figure: 3a.**
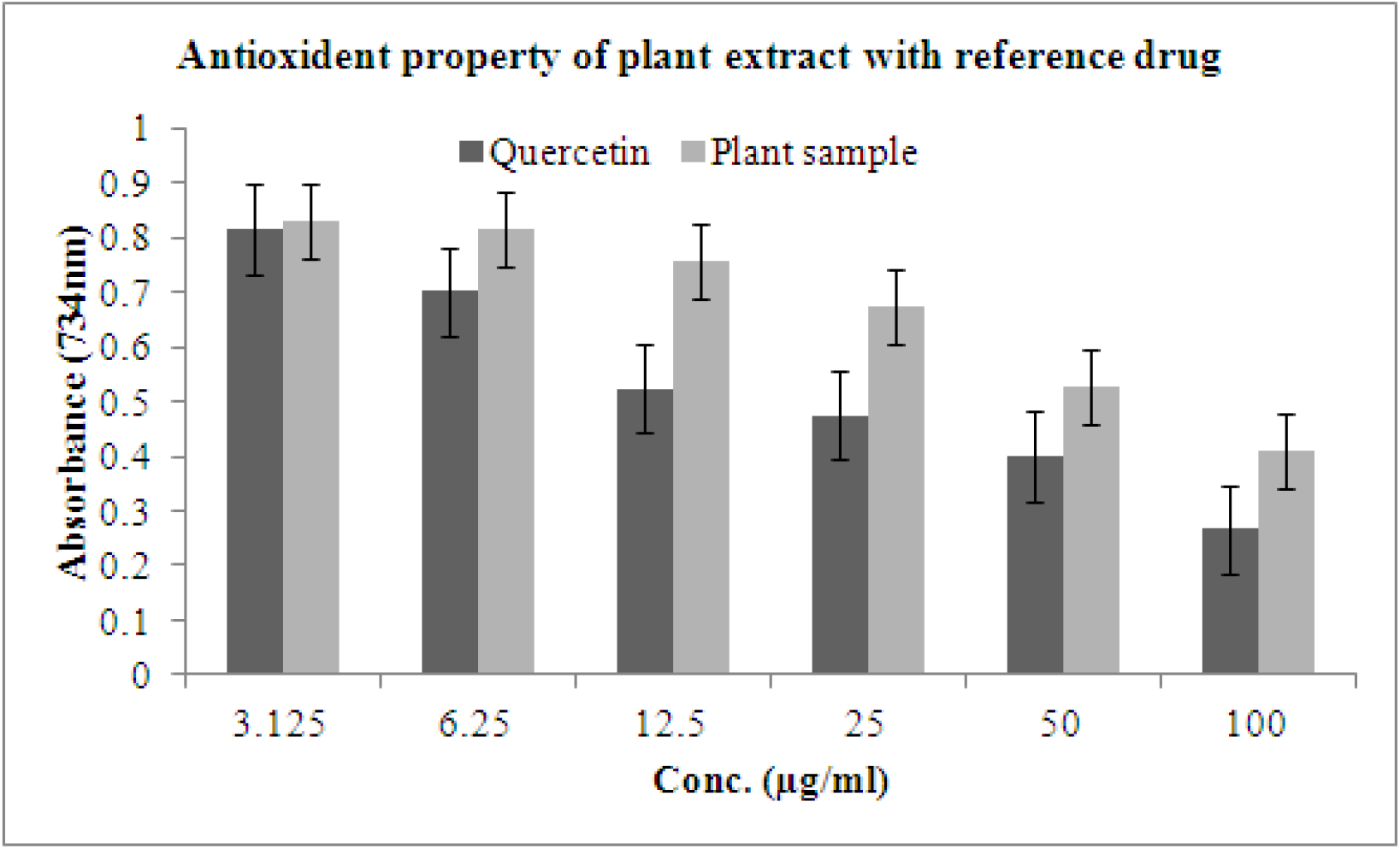
Antioxidant property prediction of *A.indicum* leaf extract with ABTS assays.

**Figure: 3b.**
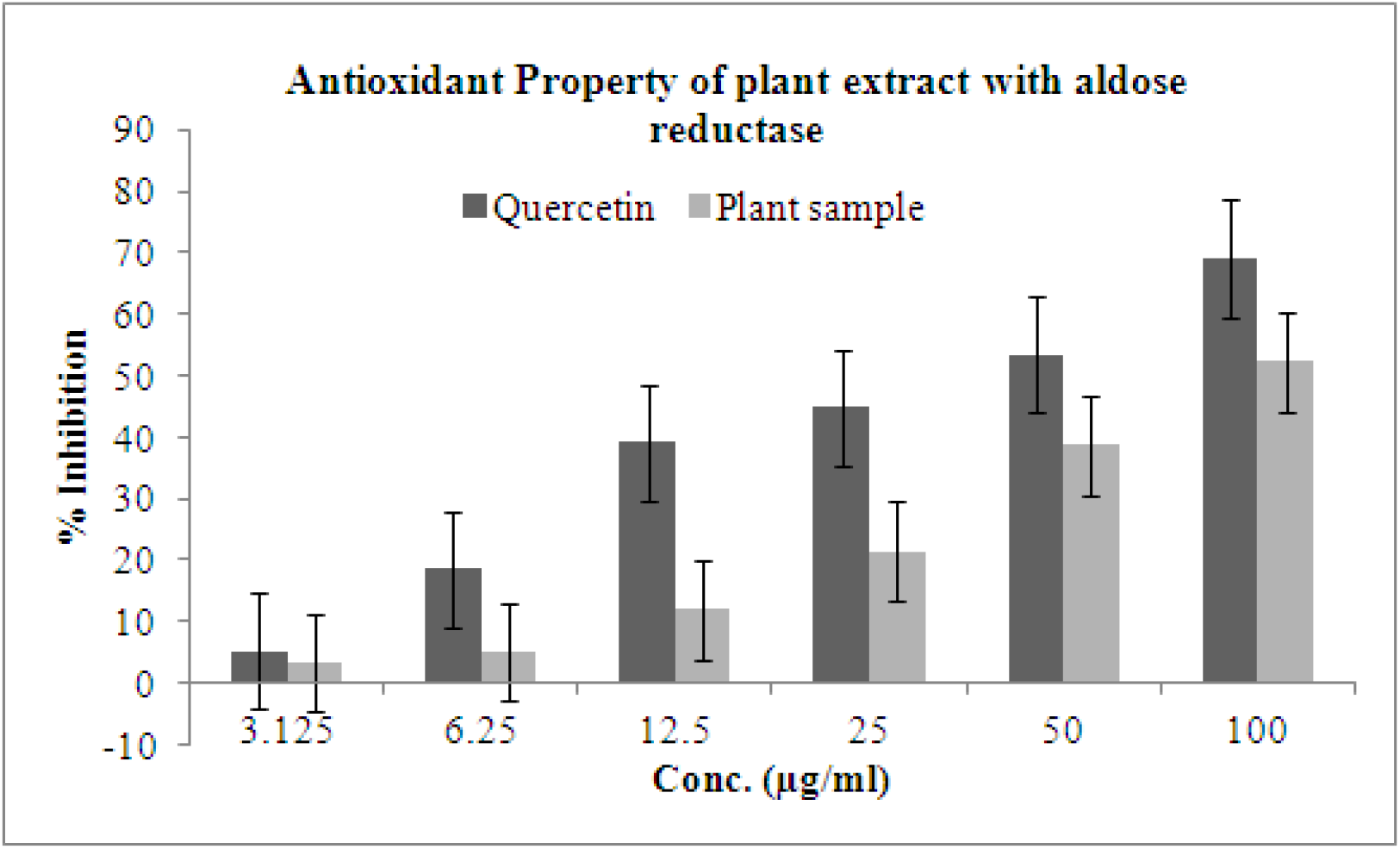
Antioxidant and inhibitory study of plant extract with aldose reductase protein.

### 3.4. Insilico analysis

The aldose reductase inhibitory and anti-oxidant properties of copper sulphate and Quercetin reference compounds along with phytochemical studies shows the strong inhibitory function. Further to screen these chemical compounds using computational drug discovery methods to design the compounds using Chemsketch v11.0 (Figure: 4) and these compounds to study drug-like prediction using pharmacophore, pharmacokinetic, molecular docking and virtual screening to predict the best target molecules. Lipinski’s “Rule of five” have applied to the designed compounds using molinspiration of default parameters such as Octonol-water partition coefficient (miLogPa <5), topological surface area (30-130), N number of atoms (<80), molecular weight of the compounds (MW <500 kDa), molecular volume (<500), hydrogen bond acceptors (n_ON_ <10), hydrogen bond donors (n_OHNH_ <5), number of rotatable bonds (<10) (Table:2). Based on the results the highlighted red color indicates the compounds were violating the drug like properties, but further to study the bioactive properties of these compounds based on the activity of receptor molecules.

**Figure: 4.**
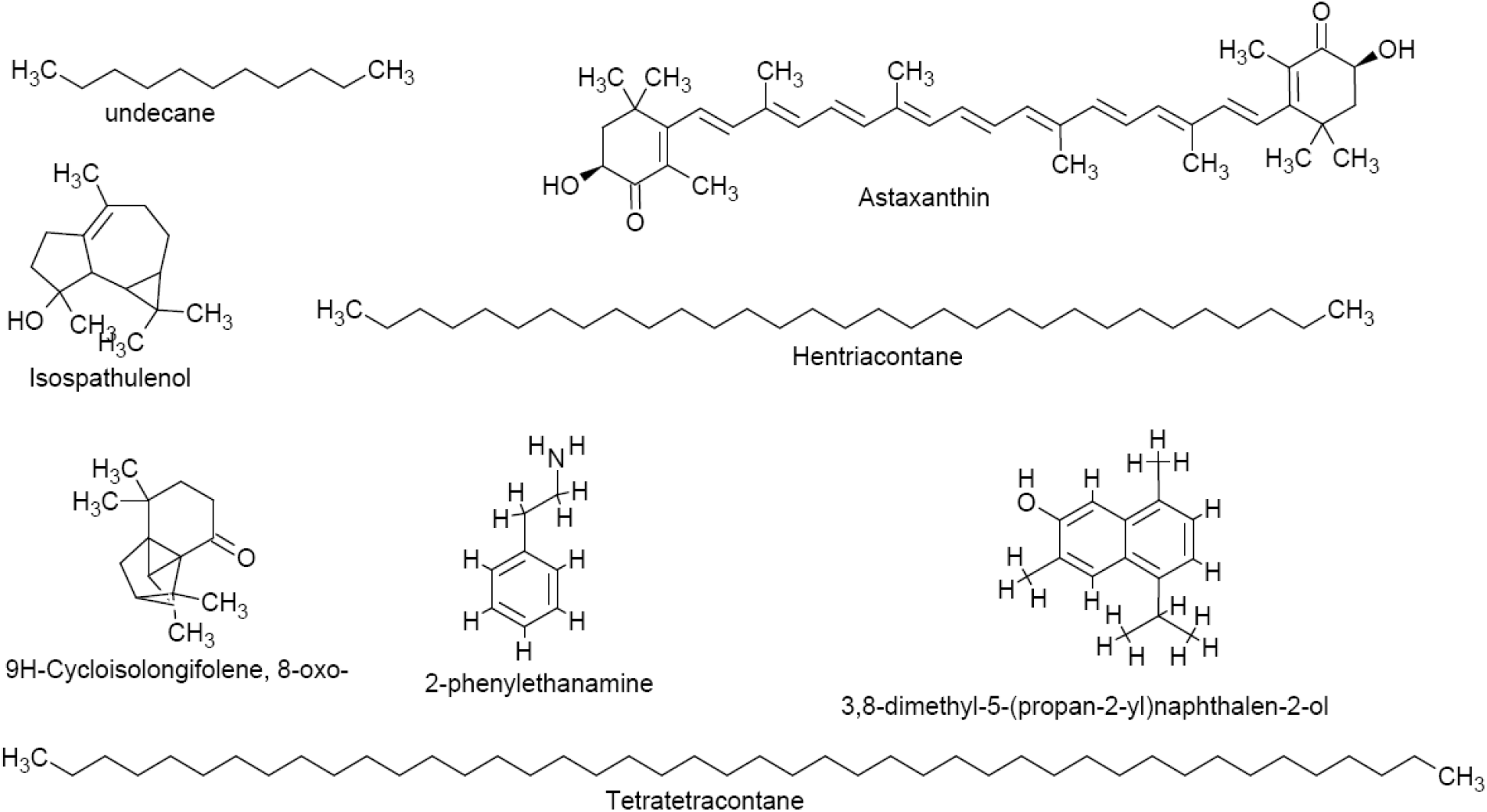
designing of phytochemicals from *A.indicum* using Chemsketch v11.0.

**Table: 2.**
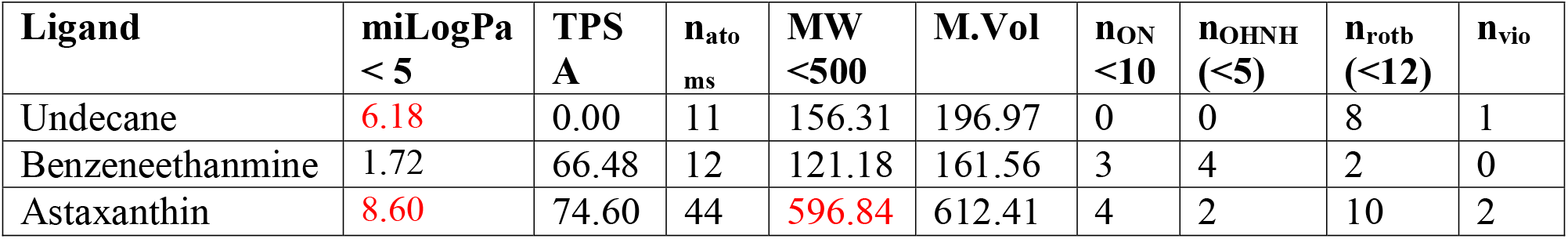
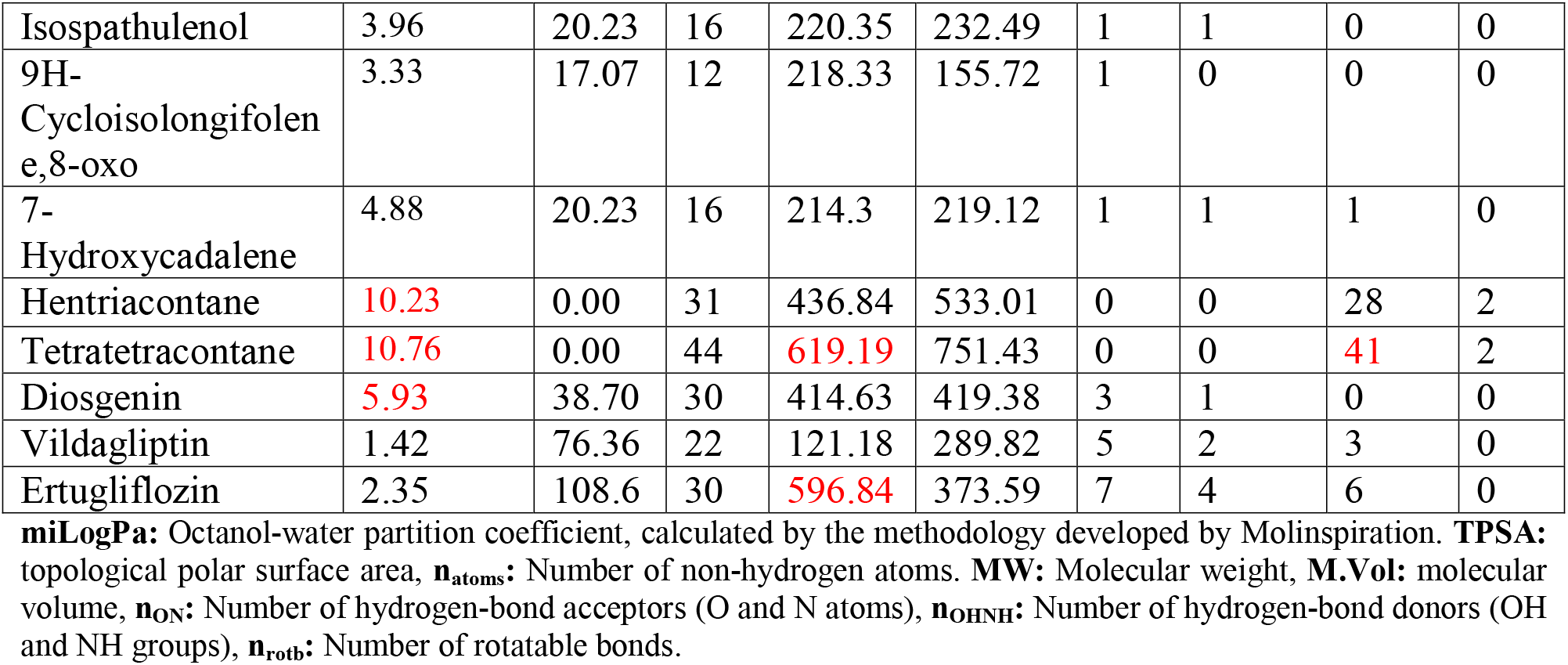
Drug-like properties of phytochemicals extracted from *A.indicum (L)*

Bioactive properties of phytochemicals are tested based on the activity scores of GPCR ligand, ion channel modulator, nuclear receptor legend, kinase inhibitor, protease inhibitor and enzyme inhibitor (Table: 3). the studies were conducted by means of online server Molinspiration to predict bioactive properties. The results are shows the yellow color highlighted the moderate functional activity with the target receptors and red color highlighted best functional activity with the target receptor.

**Table: 3.** Bioactivity of phytochemicals extracted from *A.indicum (L)*

| Ligand | GPCR ligand | Ion channel modulator | Kinase inhibitor | Nuclear receptor ligand | Protease inhibitor | Enzyme inhibitor |
| --- | --- | --- | --- | --- | --- | --- |
| Undecane | -0.84 | -0.31 | -1.08 | -0.98 | -0.98 | -0.44 |
| Benzeneethanmine | -1.71 | -1.16 | -1.95 | -2.61 | -1.85 | -1.43 |
| Astaxanthin | -0.09 | -0.62 | -0.45 | 0.16 | 0.00 | -0.05 |
| Isospathulenol | -0.78 | -0.29 | -0.81 | 0.05 | -0.39 | -0.00 |
| 9H-Cycloisolongifolene,8-oxo | -0.26 | -0.05 | -0.75 | -0.13 | -0.28 | 0.09 |
| 7-Hydroxycadalene | -0.34 | -0.21 | -0.33 | -0.04 | -0.54 | -0.06 |
| Hentriacontane | 0.03 | 0.00 | -0.03 | 0.04 | 0.03 | 0.02 |
| Tetratetracontane | -0.11 | -0.59 | -0.33 | -0.30 | -0.01 | -0.29 |
| Diosgenin | 0.05 | -0.14 | -0.57 | 0.58 | -0.06 | 0.61 |
| Vildagliptin | 0.40 | -0.04 | -0.26 | -0.19 | 1.02 | 0.17 |
| Ertugliflozin | 0.22 | 0.12 | -0.13 | 0.23 | -0.07 | 0.28 |

QSAR properties is predicted to the phytochemicals have several parameters such as surface area (Å^2^), volume (Å^3^), hydration energy kcal/mol, logP, refractivity (Å^3^), Polarizability (Å^3^), mass (amu), total energy etc. are computed for above optimized molecule using Hyperchem 7.0 Professional (table:4). The one of the QSAR parameters logP is critical parameter to express the competence of the molecule to cross the cell membrane to reach receptor site in biological systems. The logP of all compounds has positive values which represents for more hydrophilicity rather than lipophilicity.

**Table: 4.** QSAR Properties of phytochemicals identified from *A.indicum (L)*

| Ligands | SA (Å<sup>2</sup>) | SA (grid Å<sup>2</sup>) | Volume (Å<sup>3</sup>) | HE kcal/mol | logP | Refractivity (Å<sup>3</sup>) | Polarizability (Å<sup>3</sup>) | Mass (amu) |
| --- | --- | --- | --- | --- | --- | --- | --- | --- |
| Undecane | 526.95 | 460.59 | 705.23 | 5.44 | 4.86 | 52.41 | 20.96 | 156.31 |
| Benzeneethanmine | 308.47 | 333.30 | 506.09 | 15.65 | 0.88 | 38.17 | 14.00 | 157.13 |
| Astaxanthin | 747.43 | 768.36 | 1479.81 | -4.55 | 4.43 | 119.84 | 41.15 | 544.44 |
| Isospathulenol | 247.91 | 385.44 | 615.78 | 0.76 | 1.05 | 31.87 | 14.58 | 196.16 |
| 9H-Cycloisolongifolene, 8-oxo | 293.12 | 304.39 | 466.59 | 0.50 | 1.60 | 36.92 | 15.55 | 152.15 |
| 7-Hydroxycadalene | 404.91 | 437042 | 713.99 | 4.31 | 1.83 | 72.75 | 26.43 | 214.31 |
| Hentriacontane | 864.52 | 774.36 | 1546.12 | 8.59 | 12.79 | 144.43 | 57.66 | 436.85 |
| Tetratetracontane | 1430.9 | 1226.55 | 2290 | 14.58 | 17.94 | 204.25 | 81.51 | 619.20 |
| Diosgenin | 505.31 | 640.35 | 1190.57 | -2.55 | 5.32 | 120.26 | 47.39 | 414.63 |
| Vildagliptin | 550.27 | 448.98 | 753.07 | -6.60 | 1.41 | 65.75 | 25.83 | 279.21 |
| Ertugliflozin | 739.26 | 621.01 | 1037.70 | -9.68 | 6.45 | 89.27 | 33.55 | 411.69 |

### 3.5. Protein structure prediction

The aldose reductase enzyme complex with NADH-dependent oxidoreductase protein structure was selected from PDB database. The protein structure has the resolution of 2.4 Å units with the predicted global quality of 0.89± 0.05 is the average per-residue scores to estimate for a large set of models and represents the RMSD (i.e. standard deviation) between QMEAN score. VERIFY3D shows 90.51% of the residues have averaged 3D-1D score >= 0.2 (Pass) and the Overall Quality Factor A: 95.0166. Ramachandran plot was observed based on the stereo chemical energy of individual amino acids. The arrangement of amino acids with peptide bonds is scored based on torsional angles-phi (φ) and psi (Ψ)-of the residues (amino acids) contained in a peptide. Residues in most favored regions [A,B,L] 255 amino acids of 91.7%, Residues in additional allowed regions [a,b,l,p] 23 8.3%, Number of end-residues (excl. Gly and Pro) 18, Number of glycine residues (shown as triangles) 16 Number of proline residues 20 and the overall score of the amino acids allowed in the ramachandran plot shows 91.% amino acids, Based on an analysis of 118 structures of resolution of at least 2.0 Å, and R-factor no greater than 20%, a good quality model would be expected to have over 90% in the most favored regions. CastP is used to predict the active site amino acids (figure: 4). Some amino acids such as Gly18, Tyr209, Ser210, Leu212, Ser214, Pro261, Lys262 and Ser263 are used as best active site amino acids help to binds to the ligand structure.

**Figure: 4.**
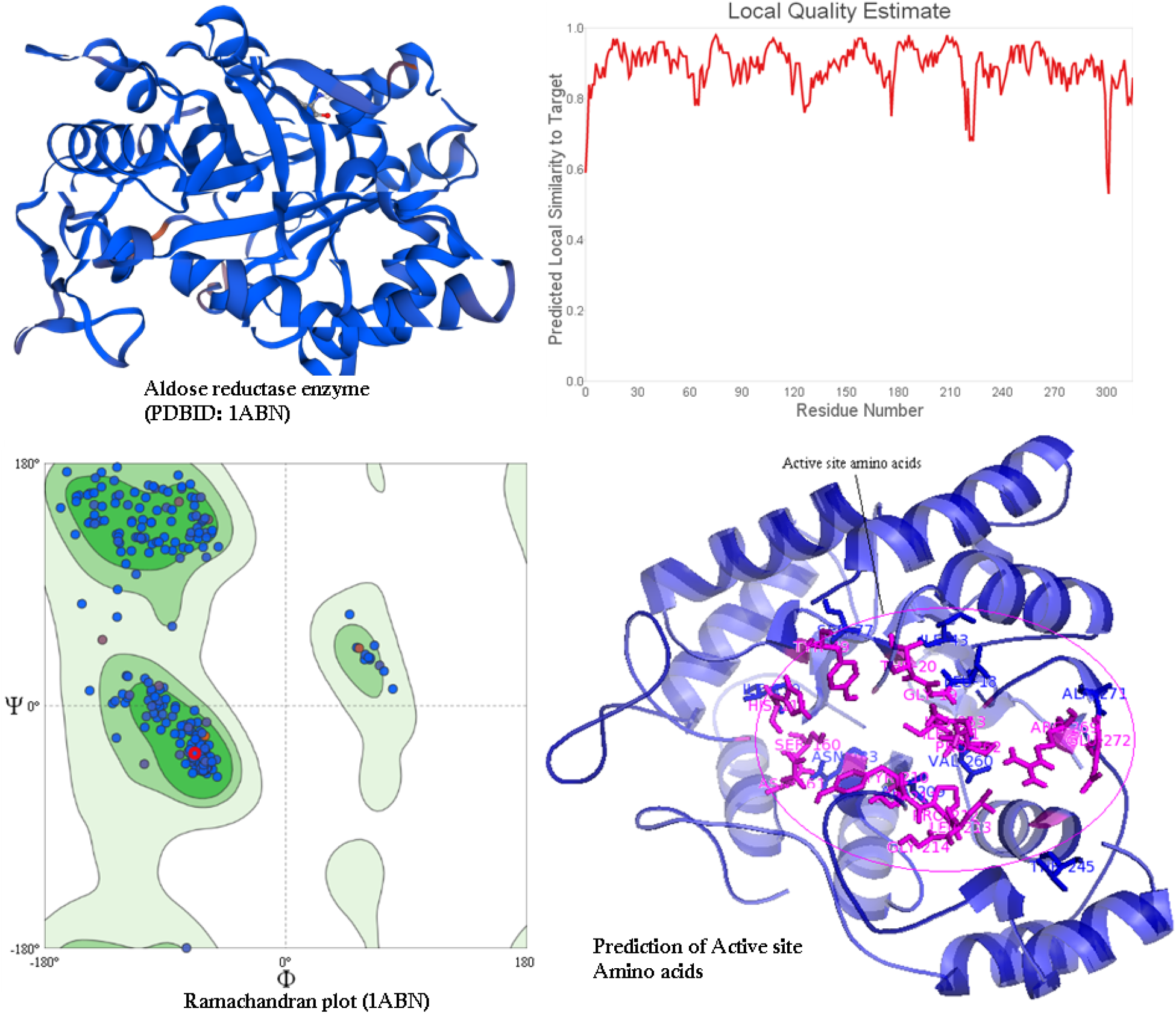
Aldose reductase protein structure, homology modeling, quality analysis and active site amino acids prediction Molecular.

Molecular Docking is performed to understand the protein-ligand interaction based on the hydrogen bonds and ligand binding energy to the target proteins. The results is compared with reference drug compound shows Ertugliflozin has 6 hydrogen bonds with the binding energy of-10.37 kcal/mol and the inhibitory constant of 2.85 nM (nanoMolar) within active site amino acids. Phytochemicals such as 7-hydroxy cadalene, Benzeneethanmine and 9H-Cycloisolongifolene,8-oxo-compounds has strong interaction with target protein by forming 3-1 hydrogen bonds with the binding energy of −7.53, −7.06 and −7.26 kcal/mol with the strong inhibitory constant of 3.04, 6.66 and 4.73μM concentration.

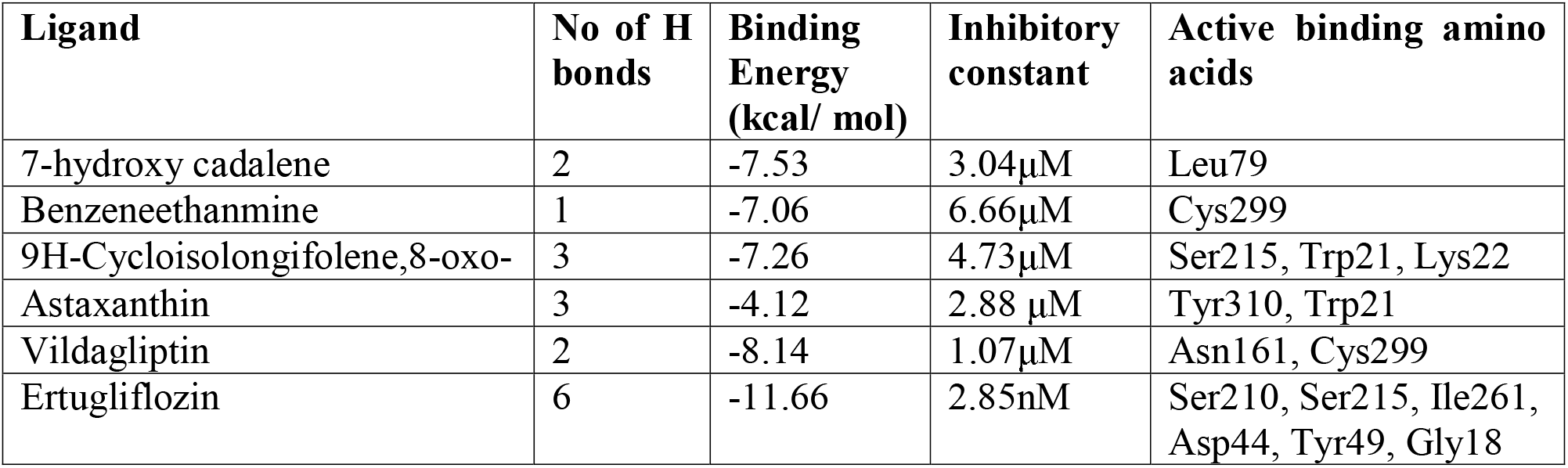

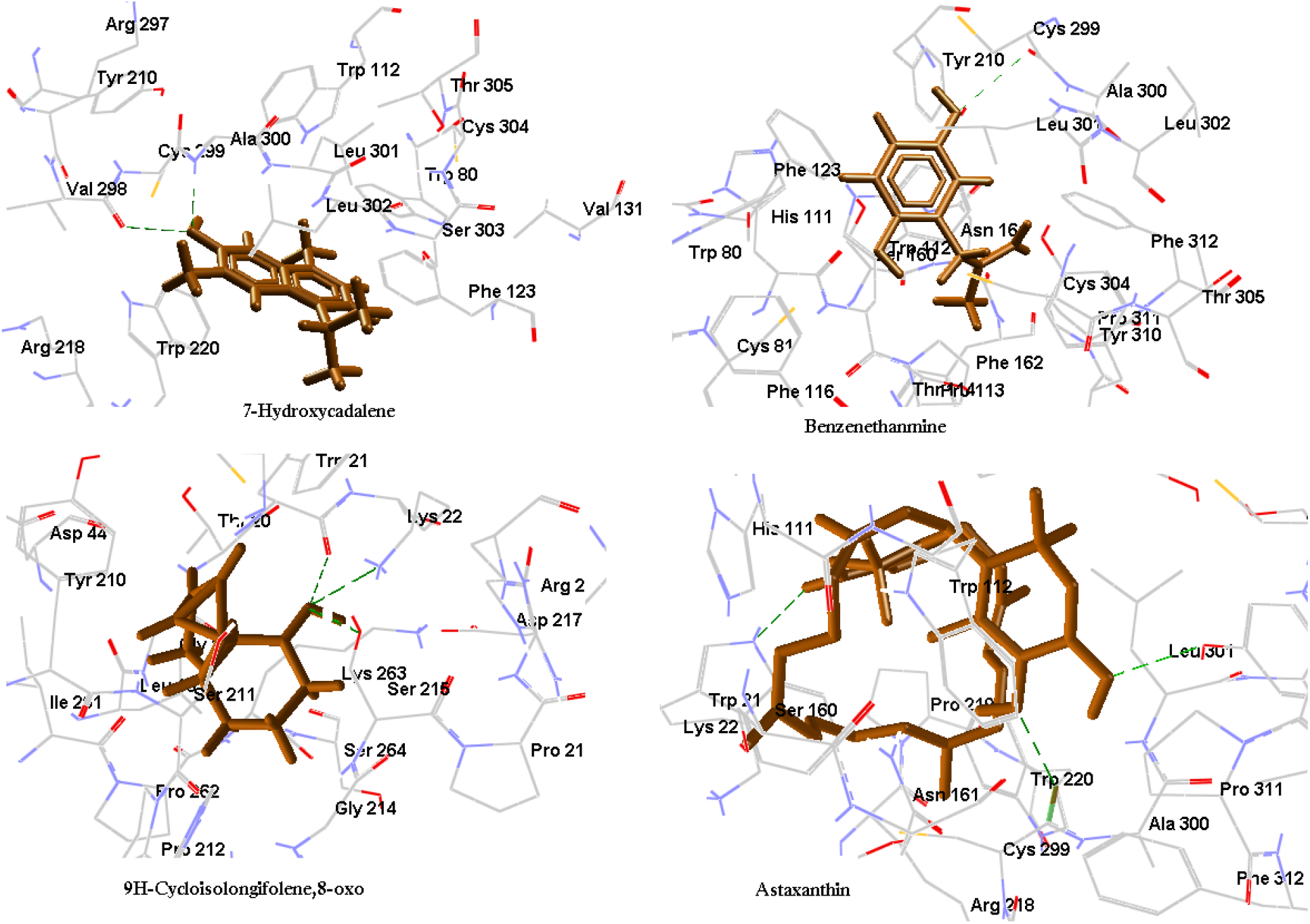

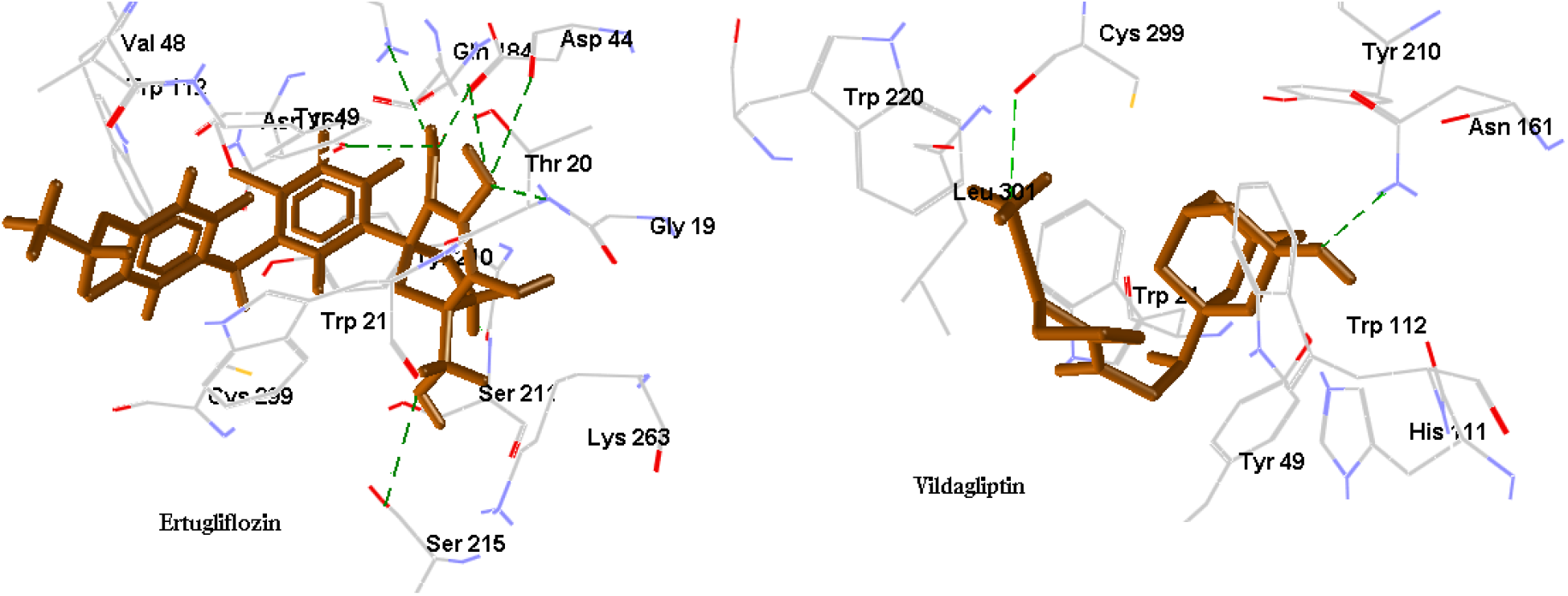

## Discussion

The current research is focused on the development of antidiabetic and antioxidant properties of the novel phytocompounds have been developed and tested for their effects. For example, Alrestatin (>2500 mg/kg) is an aldose reductase inhibitor that prevents the catalytic reaction of complicated diabetes mellitus ^[33]^. Quercetin (2.3 mg/kg) protect low density lipoprotein (LDL) from oxidation to control body glucose homeostasis in mechanism of intestinal glucose absorption and secretion of insulin to improved glucose utilization in peripheral tissues of small intestine, pancreas, adipose tissue and liver tissues ^[34]^. Copper sulphate (115-150 mg/kg) is a metal sulfate compound having copper (2+) as the metal ion; it helps to measure glucose concentration in the blood that helps to oxidoreductase activity to T2D ^[35, 36]^. Here, we focused on aldose reductase enzyme separation from rat eye ball that helps to study diabetic nephropathy. As per the literature *A. indicum* has antidiabetic activity in STZ-induced diabetic rats. The analysis of the research is based on the treatment of copper sulphate, Quercetin and phytochemicals with different concentrations. Here, we treated aldose reductase enzyme with copper sulphate (IC_50_ = 13.60μg/ml) and phytochemical (IC_50_ = 13.58μg/ml) has significant inhibition that lowers the plasma glucose level (figure: 2a, 2b). Further to study the antioxidant properties of lower concentrations, in relation to oxidizable substrates, significantly inhibit or delay oxidative processes, while often being oxidized themselves. Here, we used Quercetin (IC_50_= 1.227μg/ml) along with phytochemical sample (IC_50_ = 70.78μg/ml) has antioxidant property which is due to the presence of substituted groups such as carbonyl, phenolic, phytyl side chain, electron withdrawing group, electron donating group etc (Figure:3a, 3b). Phenolic antioxidants donate hydrogen to the radical and convert it to stable non radical product has an important role to inhibit glucose production by liver and improve carbohydrate metabolism in diabetes. Subsequently, we found that *A.indicum* Phyto-compounds have antidiabetic and antioxidant property that helps to control the diabetes.

Further have study computational drug discovery methods to screen the phytochemicals based on pharmacophore, pharmacokinetic and molecular docking study. Here, we identified 9 phytochemicals and used Lipinski’s “rule of five” for further studies. The results showed (Table: 2) that Undecane, Astaxanthin, Hentriacontane, Tetratetracontane and Diosgenin has miLogP >5 which showed the less hydrophilicity, it has less lipophilic that implied that these compounds should have less permeability across the cell membrane. The molecular weight of Astaxanthin, Tetratetracontane and Ertugliflozin has >500kda has showed less transport to the membrane that has less diffuses absorption. The hydrogen bonds and acceptors has determined based on Lipinski’s limit of <5 and <10 respectively, TPSA property helps to predict intestinal absorption, permeability and blood-brain barrier penetration that shows the screening of compounds. Further, we predict the bioactive properties were examined by calculating the activity scores of GPCR ligand, ion channel modulator, nuclear receptor legend, kinase inhibitor, protease inhibitor and enzyme inhibitor. The compounds Hentriacontane, Diosgenin, Vildagliptin and Ertugliflozin has (0.00 to 0.61 respectively) that show highly effective to enzyme inhibition, GPCR ligand, Ion channel modulator, Nuclear receptor ligand and Protease inhibitors. QSAR properties were applied to these compounds that shows all compounds is strongly accept the QSAR properties.

Further we studied the molecular docking studies of the selected phytochemicals along with reference drug compounds. Ertugliflozin is strongly bind with aldose reductase enzyme that binds to active site amino acids Ser210, Ser215, Ile261, Asp44, Tyr49 and Gly18 by forming 6 hydrogen bonds and the inhibitory constant of 2.85nM, the another reference compound such as Vildagliptin has 2 hydrogen bonds within active site amino acids of Asn161 and Cys299 with inhibitory constant of 1.07μM. These results were mapped with phytochemicals shows 9H-Cycloisolongifolene-8-oxo-and Astaxanthin has 3 hydrogen bonds interaction with inhibitory constant of 4.73μM and 2.88 μM respectively within active sites Ser215, Trp21, Lys22 and Tyr310 amino acids. Based on the docking observation, 9H-Cycloisolongifolene, 8-oxo-and Astaxanthin is strong effective function to inhibit aldose reductase enzyme and is potentially used for aldose reductase enzyme inhibitor.

## Conclusion

In summary, A.indicum leaf extract showed strong antidiabetic and antioxidant properties were observed in invitro analysis of Rat-eye ball aldose reductase enzyme inhibition. The plant extract has screened the drug-like prediction which provides good pharmacophore profiling and drug-likeness. Molecular docking results shows 9H-Cycloisolongifolene-8-oxo-and Astaxanthin compounds has strong interaction with aldose reductase inhibition and there is no toxicity in the treatment of diabetic complications. In this context, we have investigating aldose reductase inhibitors to prevent diabetic complications in animals. Further studies to isolate these selected compounds to treat aldose reductase inhibitors and cytotoxicity to predict retinopathy effect using an animal model are underway.

## Acknowledgement

The first author Lavanya L of a research scholar is performed the experiment and prepared the whole research paper. The First author is also thankful to colleagues for assisting in animal experiments and also insilico analysis. V.Veeraraghavan is a research guide to guiding experimental design, methodology preparation and result analysis. C.N.Prashantha has supported the computational drug discovery support to study pharmacophore, pharmacokinetic, molecular docking and virtual screening. Renuka Srihari provided the protocol to isolate the compounds and GC-MS analysis. We are also grateful to Department of biochemistry and Biotechnology, School of Applied Science REVA University for providing instrumentation and computational facilities.

## Conflict of Interest

The authors declare no conflict of interest.

## Abbreviations

T2DM: Type 2 Diabetes mellitus
NADPH: Nicotinamide adenine dinucleotide phosphate
GC-MS: Gas chromatography-Mass spectroscopy
NIST: National institute Standard and Technology

## Notes

### Competing Interest Statement

The authors have declared no competing interest.

